# Functional Neural Connectivity of the Mouse Brain using Photoacoustic Ultrasound Imaging

**DOI:** 10.1101/2025.03.31.645325

**Authors:** Isaac H Clark, Andrew Heinmiller, Wei Zhu, Xiao-Hong Zhu, Phoebe Strell, Zachary Roushdy, Mayuresh Vernekar, Sether Johnson, Dilmareth E. Natera-Rodriguez, Kevin Sun, Hannes Wiesner, Kelsey Haney, Wei Chen, Andrew W. Grande, Walter C. Low

## Abstract

Brain connectomes are insightful models that describe the connectivity of different regions throughout the brain. These connectomes are traditionally generated through temporal correlation of blood oxygen level dependent (BOLD) signals detected by functional magnetic resonance imaging (fMRI). Photoacoustic ultrasound (PAU) can also detect oxygenation levels while being more accessible and cost effective than fMRI. We propose the use of PAU to generate brain connectomes as an alternative to fMRI. In this study we successfully developed a pipeline for processing PAU data from whole brain scans of mice models and found that the connectomes it produced were comparable to those generated by fMRI, particularly, in detecting connections previously documented in the literature. Our findings suggest that PAU is a promising alternative to fMRI for mapping brain connectome, offering advantages in sensitivity and accessibility, making it a valuable tool for future research on brain connectivity.

## 1 Introduction

The functional connections between different areas of the brain provide the underlying basis of brain state and behavior. The network of interconnections can be visualized across the entire brain using a variety of approaches. Researchers can analyze functional neural networks through the use of brain connectomes (Bullmore, Ed, and Olaf Sporns. 2009; Fox Michael D. 2018; Bazinet, Vincent, et al. 2009). Brain connectomes represent either physical connections of axons to dendrites or functional connections of shared activity within the brain. Activity is derived from oxygen usage, which correlates to the firing of neurons. Neuronal oxygen consumption triggers largely increased blood supply to active brain regions. Thus, active areas in the brain are typically surrounded by a greater amount of oxygen rich blood (Heeger, David J., and David Ress. 2022).

Functional magnetic resonance imaging (fMRI) can detect blood oxygenation levels based on the oxygen bound or unbound state of hemoglobin. The bound form of hemoglobin, oxyhemoglobin, has no unpaired electrons, which makes it diamagnetic. The diamagnetic state has no significant effect on the magnetic resonance (MR) signal. While the unbound form of hemoglobin, deoxyhemoglobin, has four unpaired electrons, which makes it paramagnetic. The paramagnetic state leads to a decreased MR signal in regions with more deoxyhemoglobin (Ogawa, S., et al. 1990). Ultimately, oxygen rich ROIs produce a more intense signal than lower oxygen areas. Oxygen activity over time can be identified, recorded, and used to generate a brain connectome (Zerbi, Valerio, et al. 2015). However, MRI is expensive, time and skill intensive, and often not portable (Arnold, Thomas Campbell, et al. 2023). As a result, this method remains inaccessible to many researchers, labs, and clinicians.

Photoacoustic ultrasound (PAU) is much cheaper, quicker, and typically more portable than MRI machines (Joshi, Deepa, and D. S. Mehta. 2022; Wang, Lihong V., and Junjie Yao. 2016). Similar to fMRI, PAU can detect blood oxygenation. Oxygen levels are detected by using a pulsatile emission of 750 and 850 nm light, which is absorbed by hemoglobin. The pulsatile light heats and cools the molecules that absorb it, leading them to expand and contract. The expansion and contractions result in pressure waves that can be detected via the PAU. Oxyhemoglobin tends to absorb more light at 850 nm, while deoxyhemoglobin absorbs more at 750 nm. By alternating the PAU emission wavelengths to target deoxyhemoglobin or oxyhemoglobin and comparing the resulting pressure intensities, SO_2_ levels can be calculated (Li, Changhui, and Lihong V. Wang. 2009).

Prior research groups have shown that brain activity and even brain connectivity can be detected using PAU (Zhang, Pengfei, et al. 2017, Nasiriavanaki, Mohammadreza, et al. 2013). Their results showed many bilateral correlations in regions such as limbic, somatosensory, and hippocampal which have all been established in prior literature. These studies used a repetition time of 10Hz, while viewing a single coronal or axial slice. In our study, rather than using a single coronal or axial slice, we used whole brain PAU scans, developed a pipeline to map them, and generated whole brain connectomes from them. We validated the identified connections using fMRI generated brain connectomes and prior literature.

## 2 Methods

### 2.1 Animals

Female CD1 4-8 week old mice were used in this experiment. The animals were divided into three groups: group A: 3 mice used to generate the base PAU connectomes setting, group B: 3 mice used to generate the fMRI connectomes and group C: 4 mice used to conduct the four PAU connectomes under alternate PAU settings.

### 2.2 Ultrasound Acquisition

CD1 mice were anesthetized with isoflurane (1-5%), then a depilatory cream was used to remove the hair on the head of the animal. All ultrasound and PAU imaging was performed with the Vevo F2 LAZR-X Imaging System (FUJIFILM, VisualSonics, Inc., Toronto, Canada). Each animal was mounted in a stereotactic frame custom-made to work with the mouse imaging platform, which is heated to maintain body temperature and has integrated ECG and respiration monitoring. The UHF29x transducer (center frequency 21 MHz) was used in combination with a linear stepper motor to acquire coronal images of the mouse brain with matrix voxels= 0.029˂-˃0.036mm x 0.028˂-˃0.036mm x 0.5˂-˃2mm (medial/lateral, superior/inferior, anterior/posterior), and matrix dimensions= 384x(352˂-˃304)x26. The sampling rate was performed at 1/42˂-˃1/26 Hz depending on the step size varying from 0.5˂-˃2 mm respectively. Simultaneous PAU and ultra-high frequency ultrasound images were acquired with OxhyHemo Mode, which uses images taken at two wavelengths (750nm and 850nm) to calculate the oxygen saturation of the blood at each pixel and display this as a parametric map. Images were exported separately as B-mode ultrasound images and PAU OxyHemo images as a TIFF stack.

### 2.3 fMRI Acquisition

All MR experiments were performed using a horizontal 9.4 T magnet interfaced to a Varian DirectDrive console. Mice were anesthetized with isoflurane (1-5%) and secured in an animal cradle. Radio frequency transmission and signal reception were carried out using a RF surface coil with a 1.4-2.0 cm loop diameter (High Field Imaging, Minneapolis, MN). T_2_ weighted anatomical images were acquired with repetition/echo time (TR/TE) = 4000/10 ms, matrix voxels= 0.25mm x 0.25mm x 0.5mm (medial/lateral, superior/inferior, anterior/posterior), and matrix dimensions= 96×48×16. Gradient echo (GE)-echo planar imaging (EPI) based fMRI images were obtained with those same parameters.

### 2.4 Scan Mapping and Processing

Ultrasound scans were exported in the form of.tif files, each representing a single slice at a single time point. An in-house MATLAB script was created to convert the.tif files into a 4D matrix including xyz coordinates, as well as time. The resulting 4D matrix was permuted and flipped to the desired orientation, matching the Allen Brain MRI Atlas. After orientation, the matrix was saved as a.nii file and given the proper dimensional information. The same procedure was performed on the PAU data.

fMRI and MRI data were exported after acquisition as.nii files. At this point, ultrasound, PAU, MRI, and fMRI data were all in the same format and spatial orientation allowing them to be processed in the same way. With proper orientation and dimension information, copies of the Allen Brain MRI and Annotation Atlas were scaled to match the dimensions of the ultrasound and MRI scans. Tissue probability maps (TPM) were also generated from the scaled Annotation Atlas’. The fMRI and PAU images were corrected for motion using “SPM12 Software - Statistical Parametric Mapping” (SPM12) re-alignment with Quality=.9; Separation=.5; Smoothing=1; Num Passes= “Register to 1st”; Resliced Images=”All Images (1..n)”; and Masking=”Don’t Mask Images”.

The ultrasound and MRI scans were analyzed using 3D Slicer to create segmentations for all brain matter. To generate these segmentations, manual outlining of the brain tissue was used followed by using “grow from seeds” and smoothing functions. Those segmentations were then saved as binary.nii files. In MATLAB, the segmentations were then multiplied element wise to each timepoint of their corresponding scans and to the corresponding realigned fMRI/PAU scans. This stripped all non-brain material from the scan. The resulting stripped ultrasound and MRI scans were then co-registered to the Allen Brain MRI Atlas via SPM12 with separation =.1, while keeping the fMRI and PAU scans in alignment. After registration, the ultrasound and MRI images were normalized to the custom TPMs using SPM12 normalization and that transformation was applied to the PAU and fMRI scans. The parameters alternated from default were Affine Regularization=European Brains, sampling distance =.1, and bounding box and voxel sizes relevant to the scaled atlas’s. The fMRI and PAU images were then gaussian filtered using a FWHM equal to 3 times the average voxel width. Lastly, the fMRI data was temporally filtered via a highpass butterworth filter with a filter order of 2 and cutoff frequency of.01Hz.

### 2.5 Scan Connectome Generation

Prior to connectome generation, the annotation matrix was scrubbed such that certain subregions were lumped into larger regions. This ensured the same regions would be considered when comparing the ultrasound generated connectome with an MRI-generated connectome. Additionally, this method could be used to ensure regions smaller than the voxel size would not be considered. With both the ultrasound and photoacoustic data registered to the Allen Brain Atlas, the oxygenation signal was then averaged for each respective region represented in the scrubbed annotation matrix. Those region-averaged signals were then correlated against each other within the annotation matrix to produce an adjacency matrix.

### 2.6 Ultrasound Connectome Accuracy Validation

To validate the accuracy of the ultrasound brain connectome generated, we compared it to a traditional fMRI generated connectome as well as conducted a search of the scientific literature regarding reports of anatomically and functionally identified neuronal connections. A comparison to the traditional fMRI-generated connectome was performed by calculation of the mean squared error for all correlations. This was calculated for each connectome generated. Literature analysis was performed only on the averaged connectome generated from 3 mice scans with step sizes of 0.5mm and a sampling rate of 1/42 seconds.

### 2.7 Scientific Literature Analysis

To further investigate the validity of the PAU connectome, ROIs with a signal correlation value greater than 0.6 were identified and searched for physical or functional connectivity. Physical connections were identified using the Allen Connectivity Atlas. Pubmed searches were conducted in order to determine if the suspected connected ROIs shared a previously established functional network.

## 3 Results

### 3.1 Mapping & Filtering

To ensure the scans occupied the same 3D space, all scans were mapped to the Allen Brain Atlas. Four types of scans were acquired, two structural scans including MRI and ultrasound, and two functional scans including fMRI and PAU. These scans were successfully mapped to the Allen Brain Atlas (Figure 1). The PAU scan acquired in the initial study had a finer spatial resolution than the fMRI scan with a voxel size of 0.0004075 mm^3^ compared to 0.03125 mm^3^, yet there is a clear distinction between structural image detail in the PAU scan compared to the fMRI scan. These structural image detail differences can be attributed to differences in the point spread function, noise to signal ratio, and signal sensitivity due to alternate acquisition methods (Chen, Chuan, et al 2020). PAU images acquired at greater depths and/or with structures between the points of interest and the transducer (such as the skull), will alter the point spread function and the noise to signal ratio can increase.

**Figure 1:**
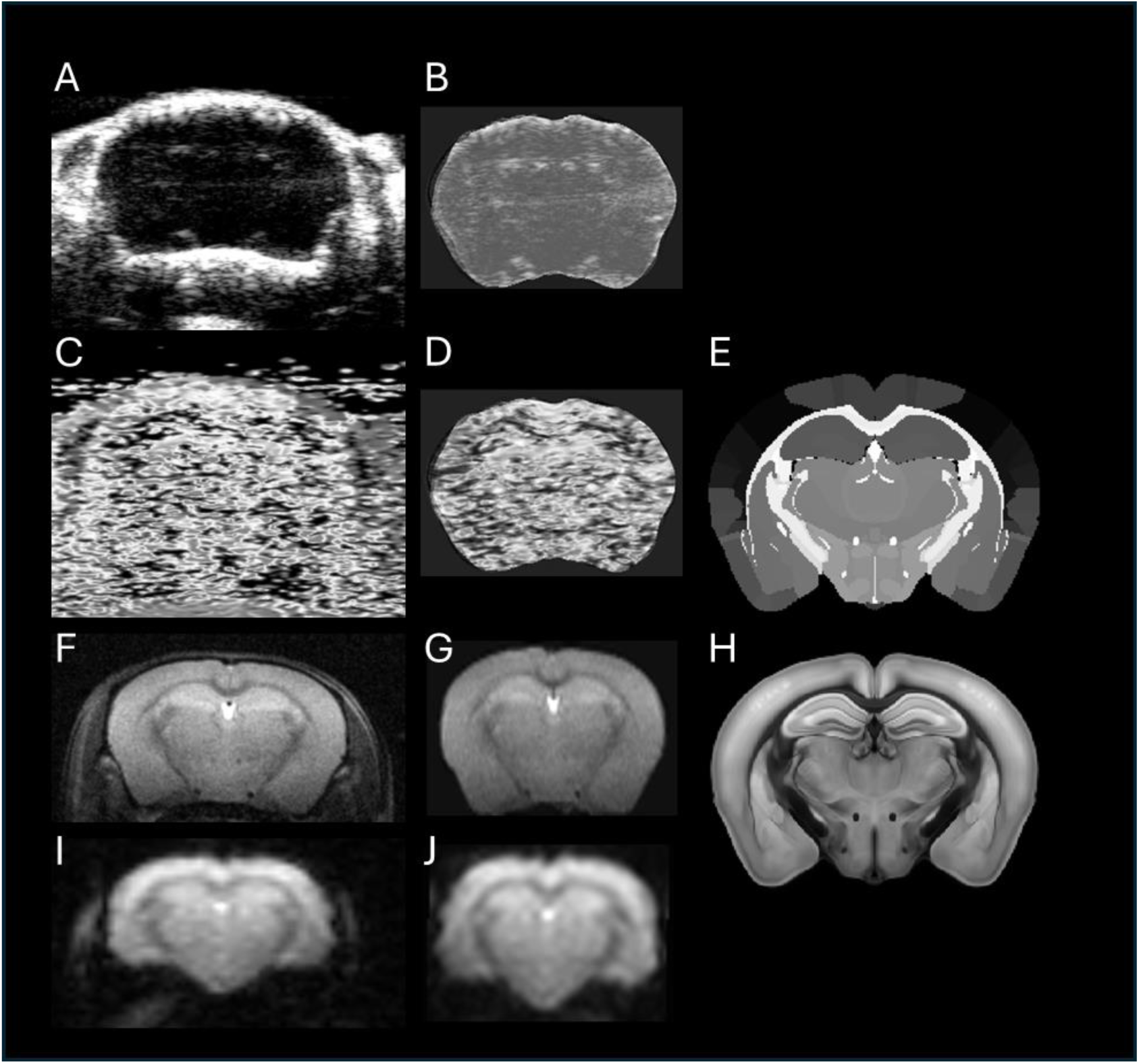
A-B: Ultrasound Scan Pre and Post Mapping. (Animal_ID:02-18-17-00-03). C-D: PAU scan pre mapping and post mapping. E: Allen Brain Annotation Atlas. F-G: T2 MRI scan pre mapping and post mapping (Animal_ID:run-29). H: Allen Brain MRI Atlas. I-J: fMRI scan pre and post mapping

Additionally, due to PAU’s simultaneous acquisition of deoxy and oxy hemoglobin, it has the potential of greater sensitivity which may explain its focus on image contrast over structural detail. In the ultrasound, fMRI and MRI scans, there is a clear distinction between the cranial cavity and non-brain matter. However, this is not true of the PAU scan. We can be certain though that the stripped and mapped PAU scan reflects the blood oxygenation signals within the cranial cavity as it was aligned with the ultrasound scan during stripping. The normalized ultrasound scan is significantly stretched/compressed to fit the atlas, which indicates scaling issues in the original scan and/or anatomical differences between the ultrasound scan and the atlas. Mild stretching/compressing is also observed in the normalized MRI and fMRI scans, indicating those same issues to a lesser extent. These issues are intrinsic to the normalization process, which ensures the scans are in alignment despite scaling or anatomical differences.

To explore a variety of processing parameters, data sets were filtered with different sampling rates, sample sizes, as well as with and without temporal smoothing. The fMRI and PAU data both had standard motion correction applied through SPM realignment and gaussian smoothing with full width at half maximum equal to three times the average voxel length. Temporal smoothing was applied to the fMRI data through a high pass butterworth filter with cutoff.01 Hz. Due to the sampling rate and temporal sample size, the butterworth filter could not be applied to the PAU data and result in viable data.

The original sampling rate of the PAU scans were 1/42 Hz in which a total of 8 time samples were acquired while the fMRI scans were acquired at 1 Hz with a total of 300 samples for one imaging run. We decided to compare samples with the same parameters, and thus a copy of the original fMRI data was downsampled from 1Hz to 1/42 Hz and only the first 8 data points of that downsampled data were kept.

### 3.2 Alternate Filtered Connectomes

The downsampled, non-temporally filtered, and temporally filtered fMRI data as well as the PAU data all had adjacency correlation matrices generated from them based on the ROIs present in the fMRI area scanned limited by the fMRI surface coil. Those adjacency matrices were averaged accounting for the 3 scanned mice and the correlation matrices were then compared (Figure 2). The correlation matrix generated from the fully filtered fRMI data was presumed to be the most accurate and most devoid of false positives, therefore acting as the control for the PAU generate connectome. We see significantly more functional connectivity in the PAU generated connectome with 4046 connections (r=having a correlation value above 0.5) compared to the fully filtered fMRI connectome only having 62 connections (r=with a correlation value above 0.5). The mean squared error (MSE) between the PAU and fMRI connectome was 0.0444. This result suggests either many additional false positives in the PAU connectome or a greater sensitivity of PAU allowing for identification of connections unidentifiable through fMRI.

**Figure 2:**
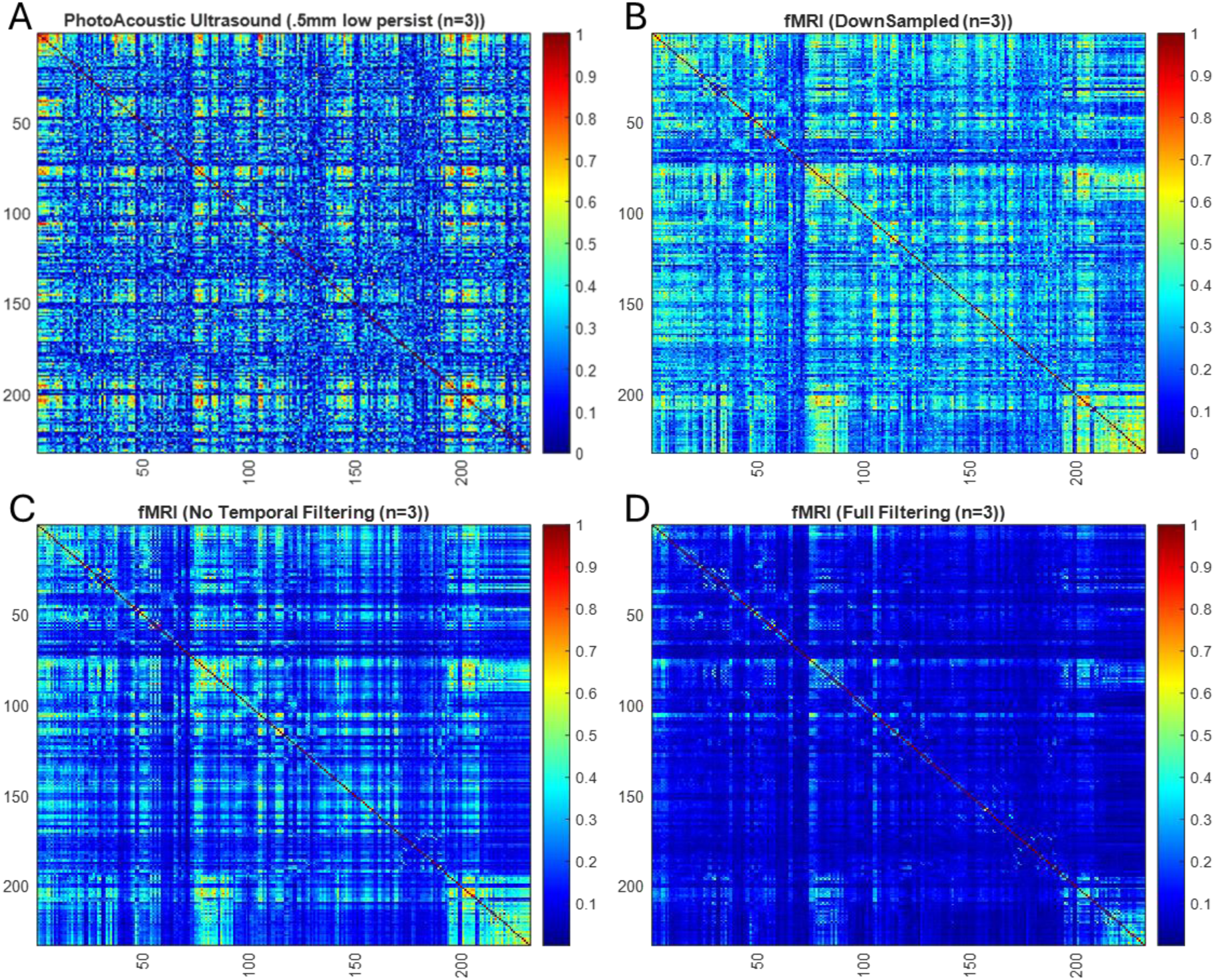
Adjacency Connectome Matrices generated from alternated scans and processing methods. A: PhotoAcoustic Ultrasound generated Connectome. B: DownSampled fMRI generated connectome. C: Non-Temporally filtered fMRI generated Connectome. D: Fully Filtered fMRI generated Connectome.

Due to the temporal sample size, the PAU data could not be temporally filtered. As such we expected it to share greater similarities with non-temporally filtered data. When PAU data was compared to non-temporally filtered data, a smaller MSE of.0306 was found. As the PAU data additionally had a lower sampling rate and smaller temporal sample size, we would expect fMRI data with the same parameters to generate a connectome most similar to the PAU data. However, this is not the case as the resulting connectome had a MSE of.0356. This implies that when downsampled the fMRI data may have some artifacts introduced or the inherent differences between oxygen detection from fMRI and PAU are more apparent at this sample size.

### 3.3 Regions of Interest

The connectomes generated by PAU and fMRI reveal select ROIs with similar connectivity patterns. For instance: the primary motor area (right), Ammon’s horn (right), and the periaqueductal gray (right). These connectivity patterns for the specific ROIs are shown in Figure 3. When analyzing these specific ROIs, we found that all connectomes identified connectivity between specific ROIs such as the motor cortex and ammon’s horn. This supports the known functional connection between these ROIs through the somatomotor network. Of note, the fully filtered fMRI scan showed the least number of connections of the fMRI generated connectomes, indicating that the other fMRI variations may have many false positives. The PAU generated connectome also showed many additional connections that the fMRI did not. Whether these connections are true or not was further evaluated in our literature analysis.

**Figure 3:**
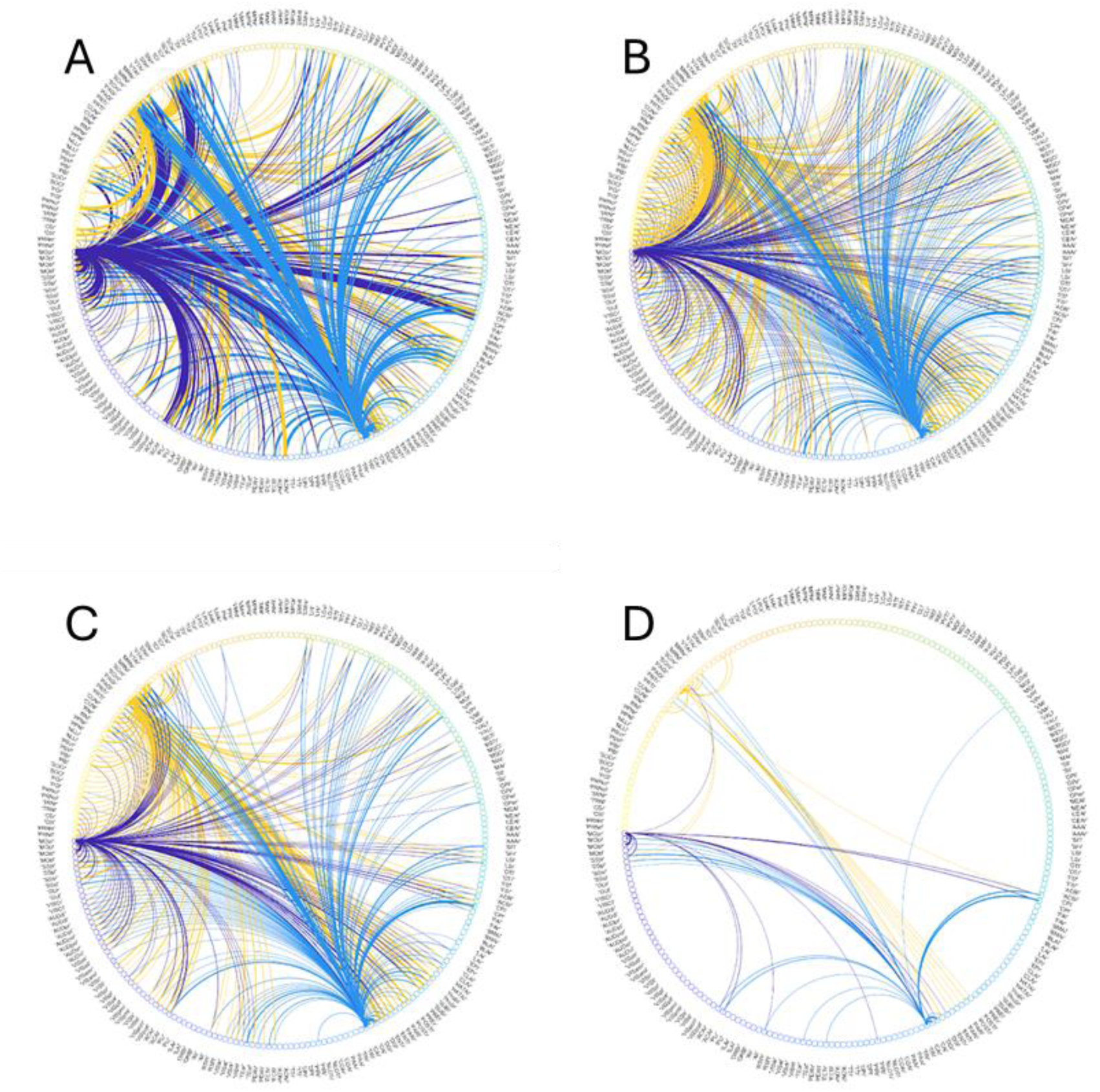
Circular Graphs of Areas with High Connectivity (ROIs: Primary motor area: Purple, Ammon’s horn: Blue, Periaqueductal gray: Yellow). All graphs show connectivity between the Primary motor area and the Ammon’s horn. A: PAU generated circular graph. B: DownSampled fMRI generated circular graph. C: Non-Temporally filtered fMRI generated circular graph. D: Fully Filtered fMRI generated circular graph.

### 3.4 Full Connectome Range

One limitation often present in fMRI scans is physical range. The fMRI data acquired in this study made use of an RF surface coil in this study to properly transmit and receive the radio waves, which limited the data acquisition area covering the brain. The PAU scan did not have this limitation, and therefore was able to acquire oxygenation signals throughout the entire brain (Figure 4). The analysis of these additional ROIs provides the opportunity to further understand the networks throughout the brain and their extent. For the purposes of this paper, the validation of these additional ROIs was saved for future research.

**Figure 4:**
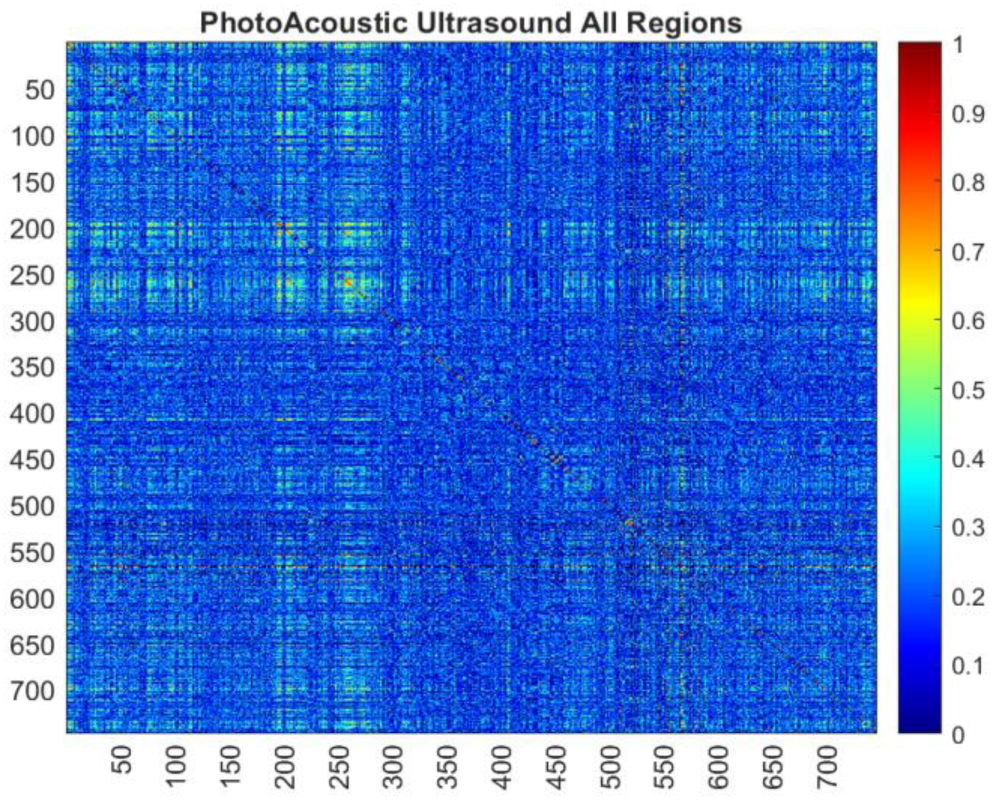
Expanded PhotoAcoustic Ultrasound Connectome.

### 3.5 Alternate PAU Acquisition Parameters

The original parameters of the PAU had a step size of.5mm, sample rate of 1/42 seconds, wavelengths emitted 750nm & 850nm, and low frame averaging (persists). To generate the most accurate PAU connectome, we adjusted these parameters and assessed the resulting errors in comparison with the connectome generated from the fully filtered fMRI scan (Figure 5). Low persistence decreased the MSE more than no persistence with the largest difference being present in the.5mm step size scans. By changing the step size, there was a noted sizable effect on the MSE with the lowest MSE being present on.5mm step size scans. When a wavelength of 850 nm was used rather than 750 and 850 nm, the resulting error was greater. The averaged connectome had the smallest MSE, which is representative of a larger sample size improving the accuracy and reliability of the results.

**Figure 5:**
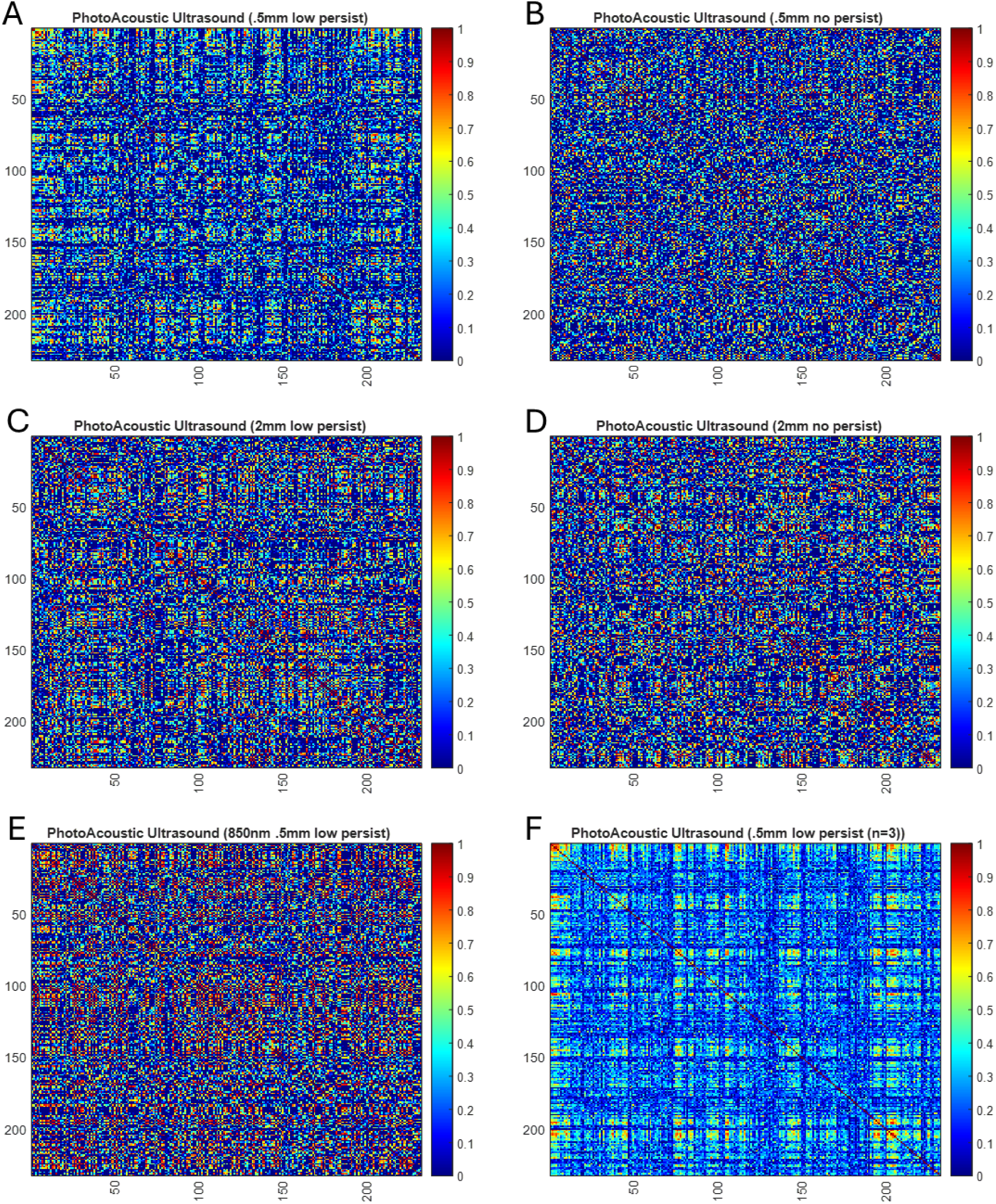
PAU Generated Connectome with Alternate Acquisition Parameters. A: 0.5mm step size with low persists. B: 0.5mm step size with no persist. C: 2mm step size with low persists. D: 2mm step size with no persists. E: 0.5mm step size with low persists and only 850nm wavelength emission. F: Average of 3 samples with 0.5mm step size with low persists.

### 3.6 Scientific Literature Analysis

To assist in ruling out false positives, we conducted a literature review of the more strongly connected PAU ROIs with correlation.6 or greater in Table 1. This value was chosen as it allowed us to focus on the highest ∼3% of correlation coefficients in the PAU connectome, increasing the confidence of a true connection. Firstly, it was recorded whether the corresponding fMRI correlation coefficient was.2 or greater. This value was chosen, as similarly, it corresponded to the highest ∼3% of correlation coefficients in the fMRI connectome. We searched through the Allen Connectivity Atlas to determine if direct connections existed between the suspected ROIs. Additionally, we identified which networks, circuits, loops, or systems the seed and suspected ROIs were associated with and if they both shared any in common. Approximately 85% of the PAU suspected connections had either a corresponding fMRI correlation value greater than.2, projections connecting or passing directly between them according to the Allen Connectivity Atlas, and/or literary research showing a shared functional connection via a shared network. The remaining ∼15% of these strong connections were highlighted (in blue) as either potential false positives or connections that PAU had an increased sensitivity and ability to detect that should be further investigated in future research.

**Table 1:**
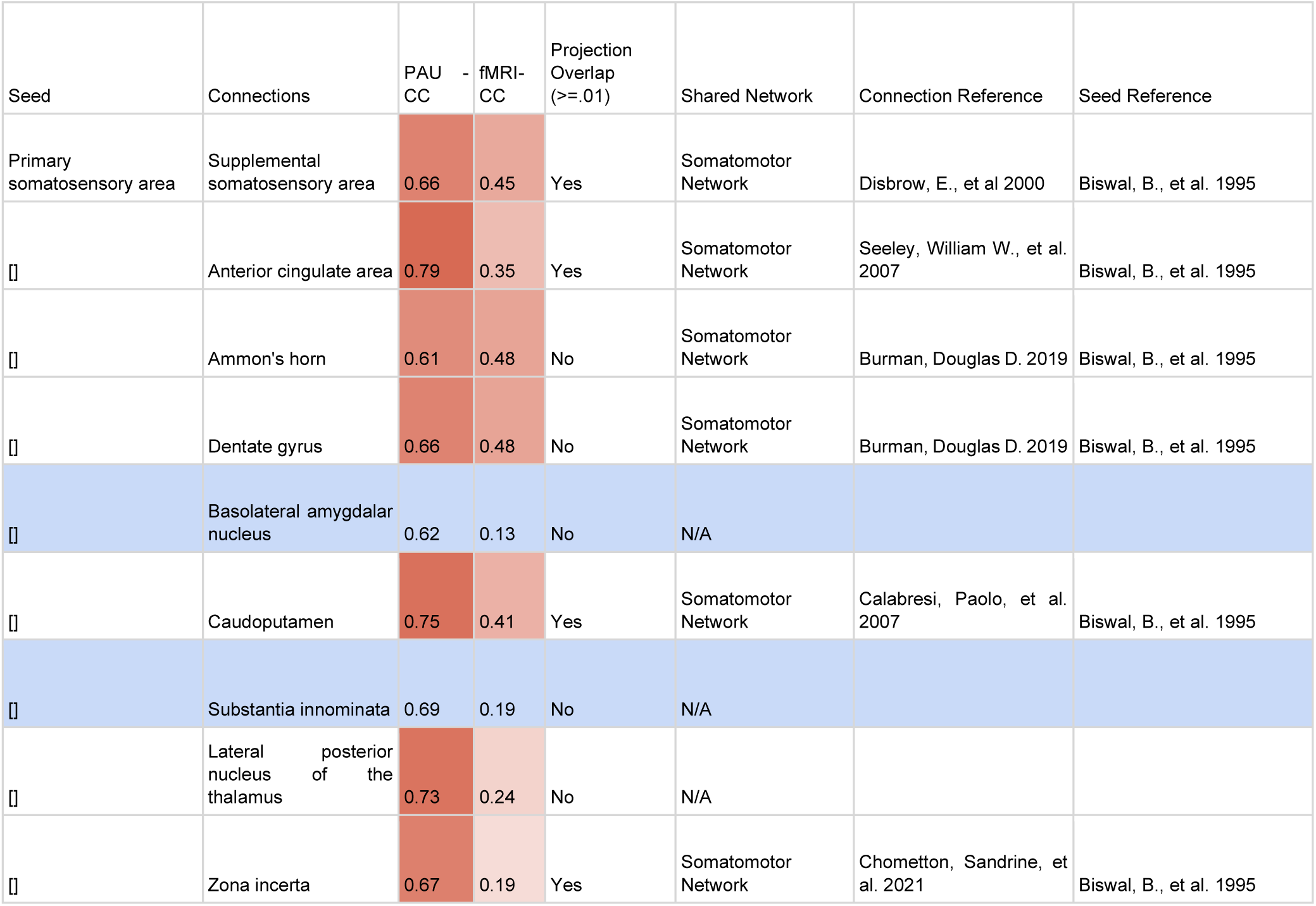
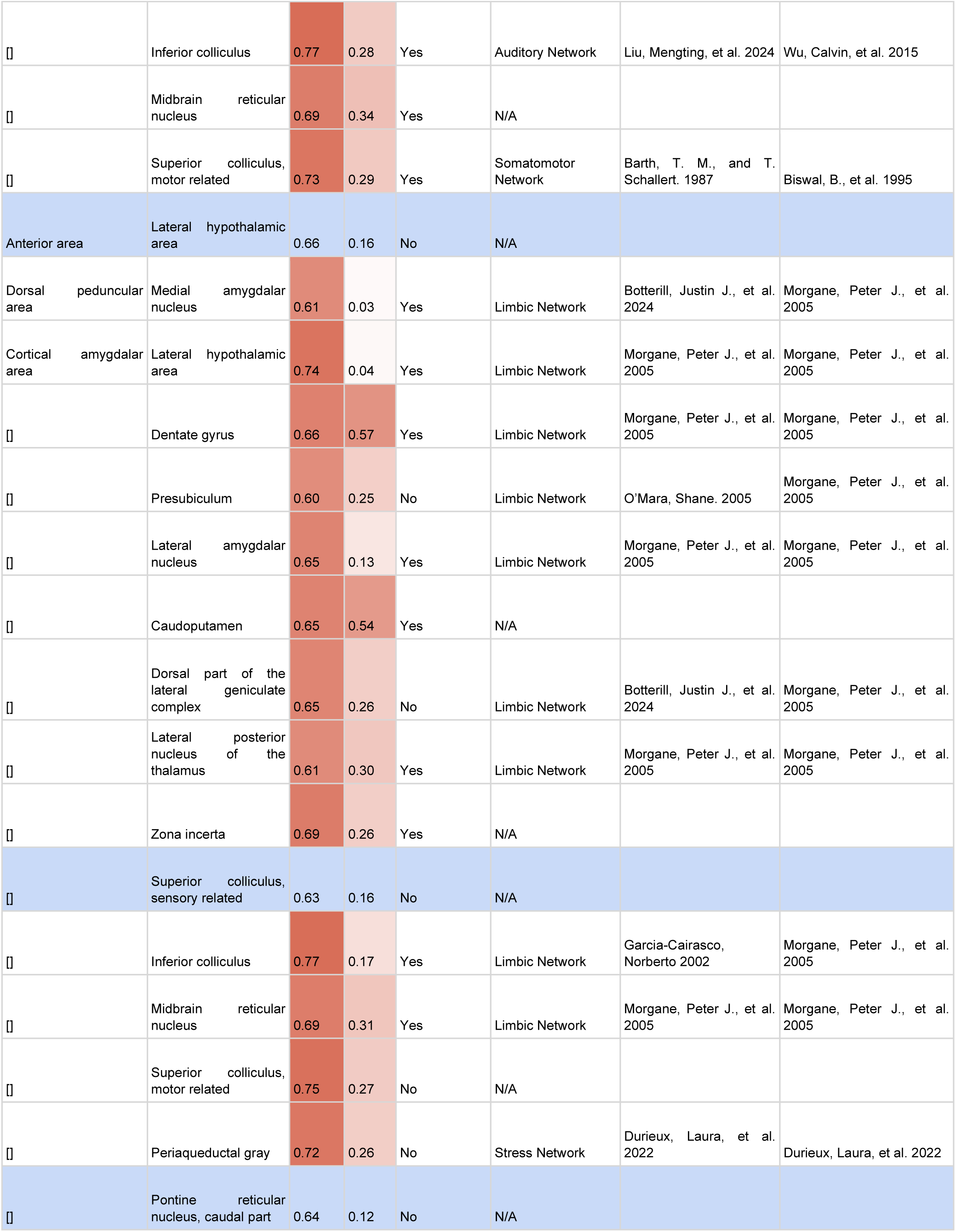
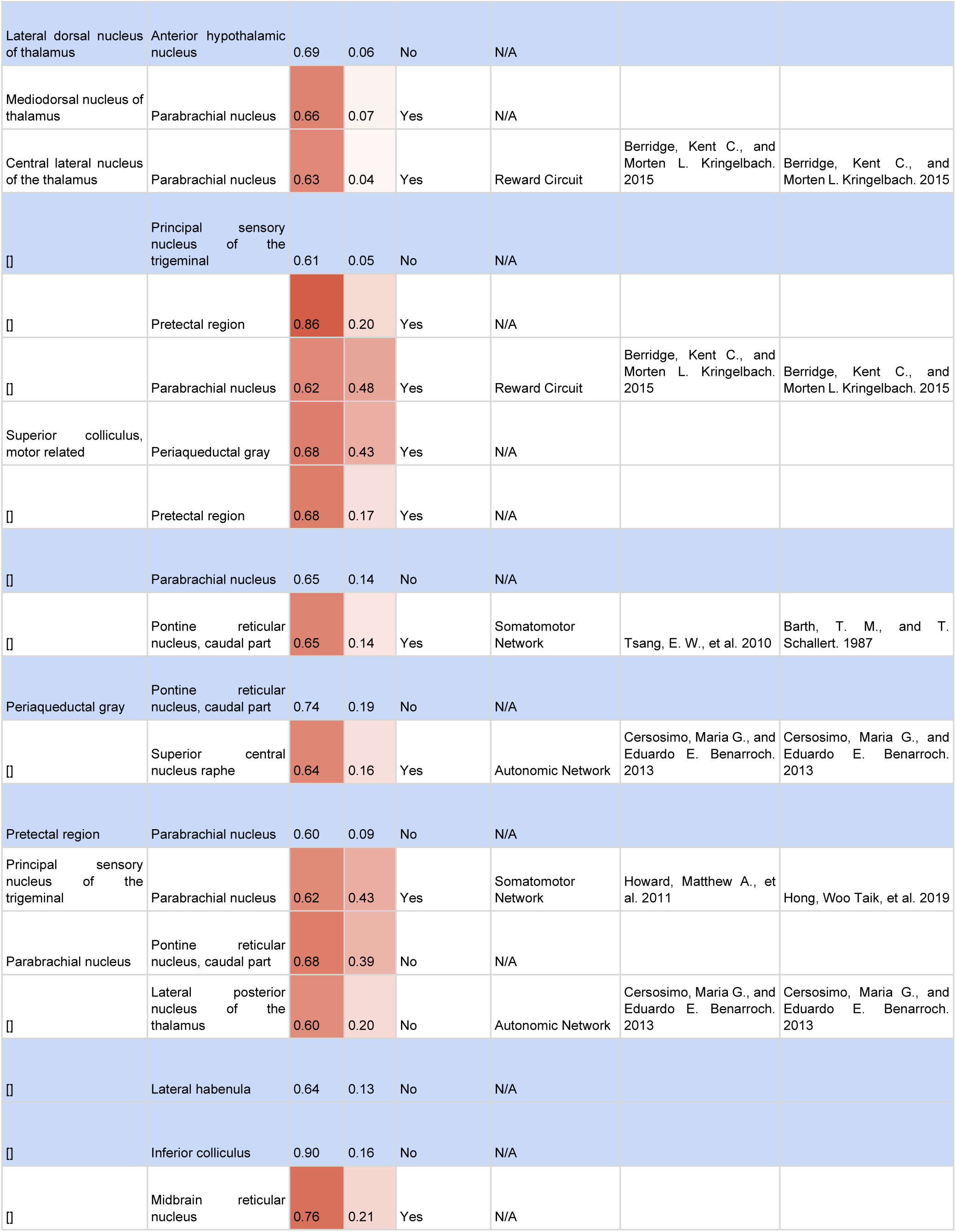
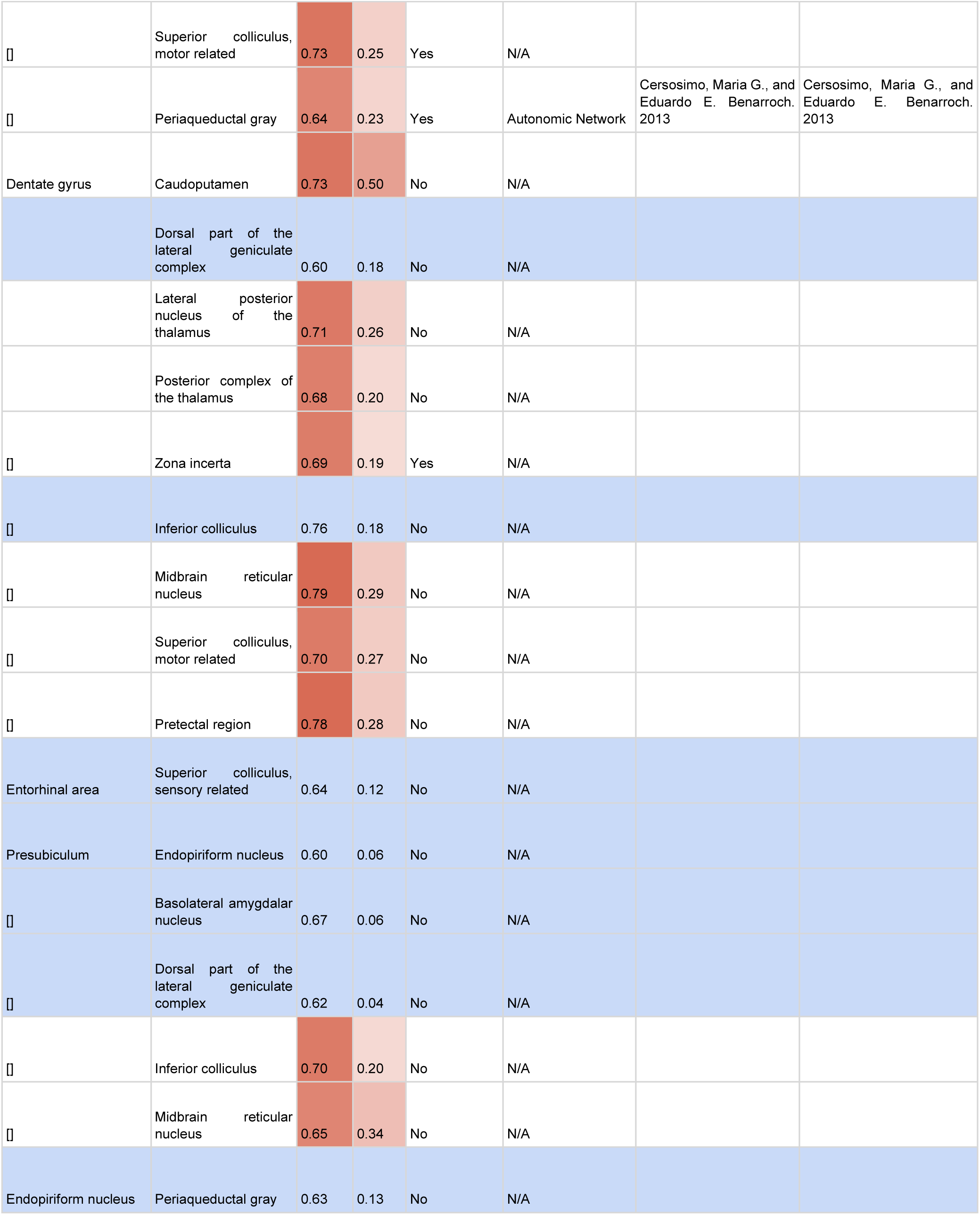
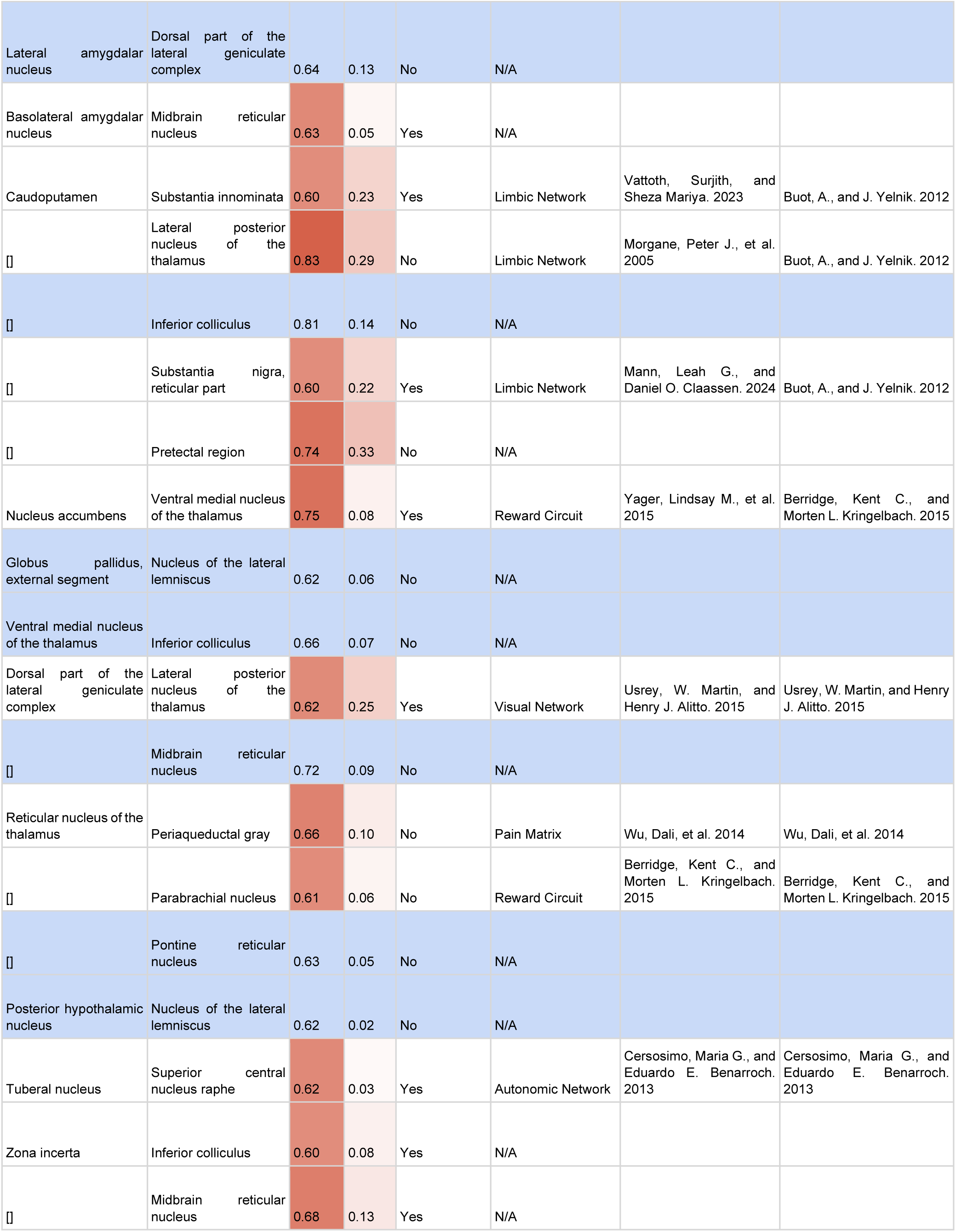
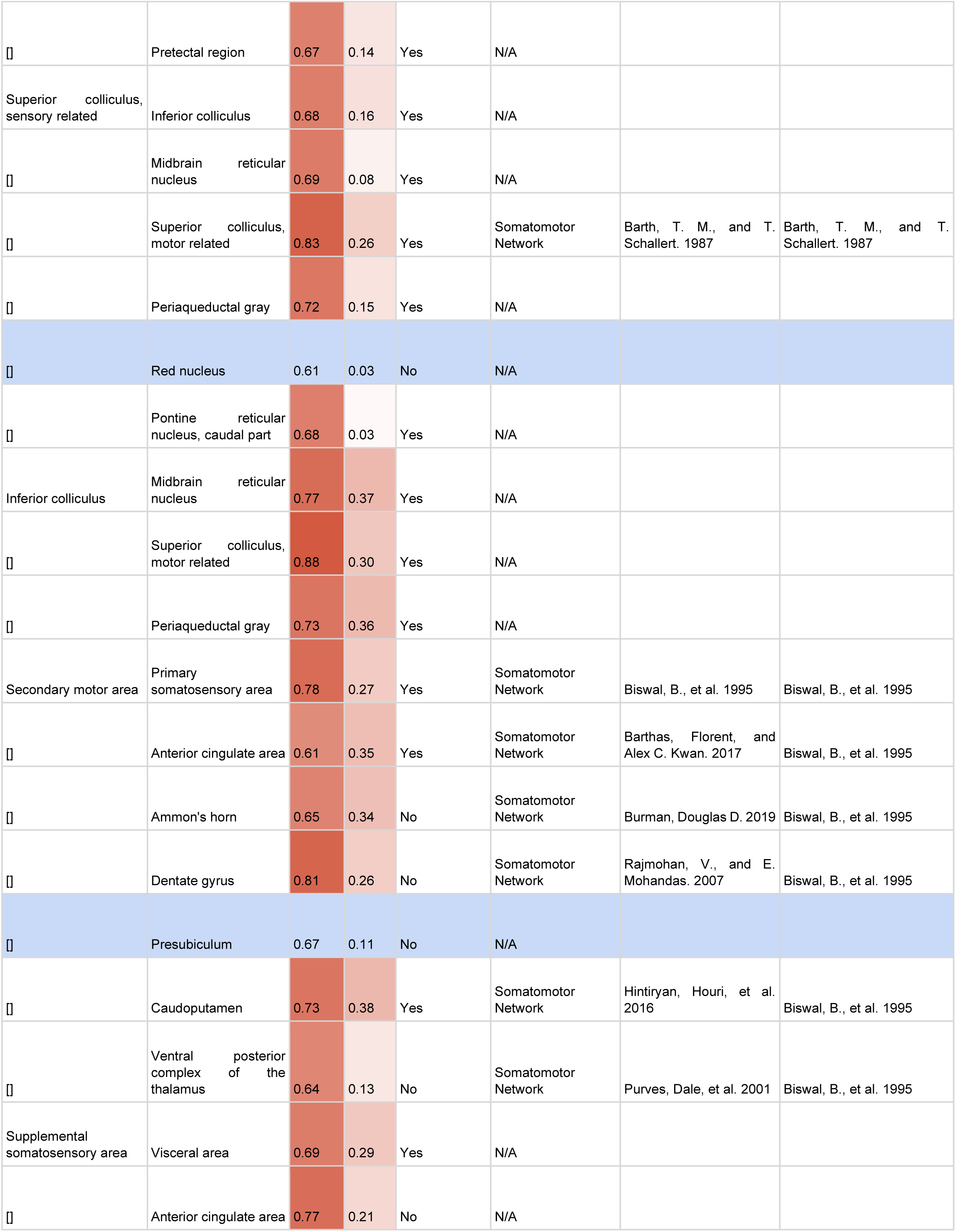
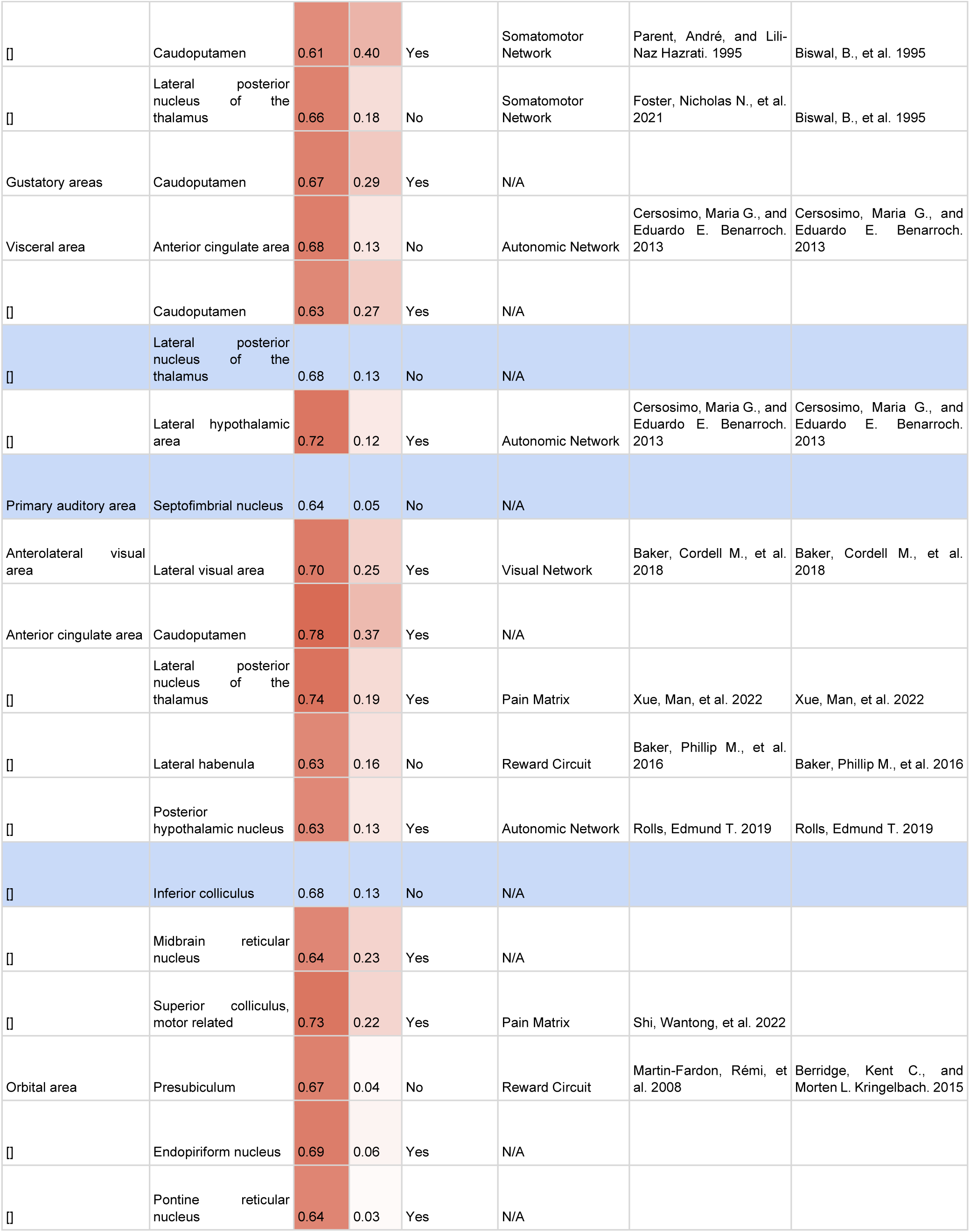
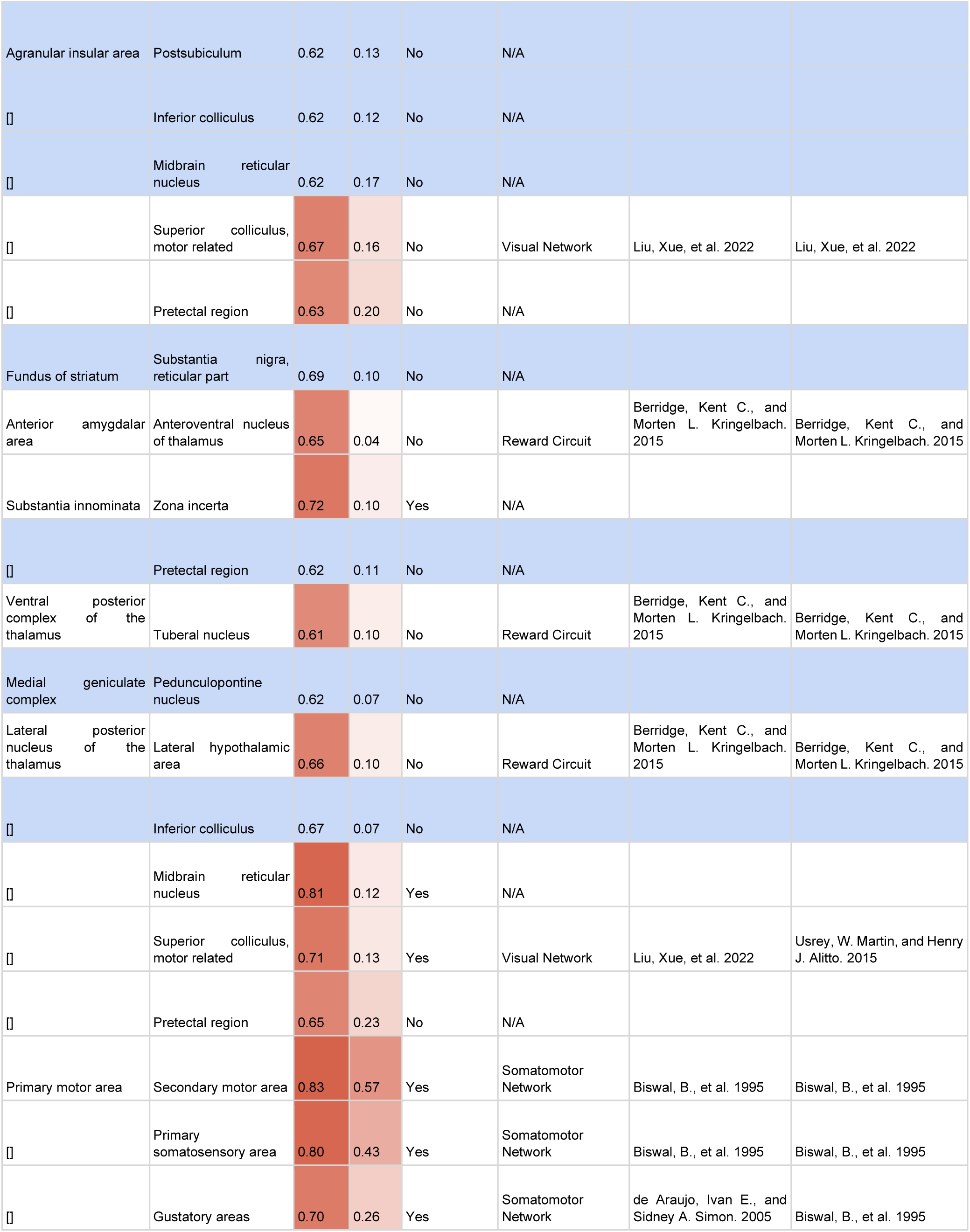
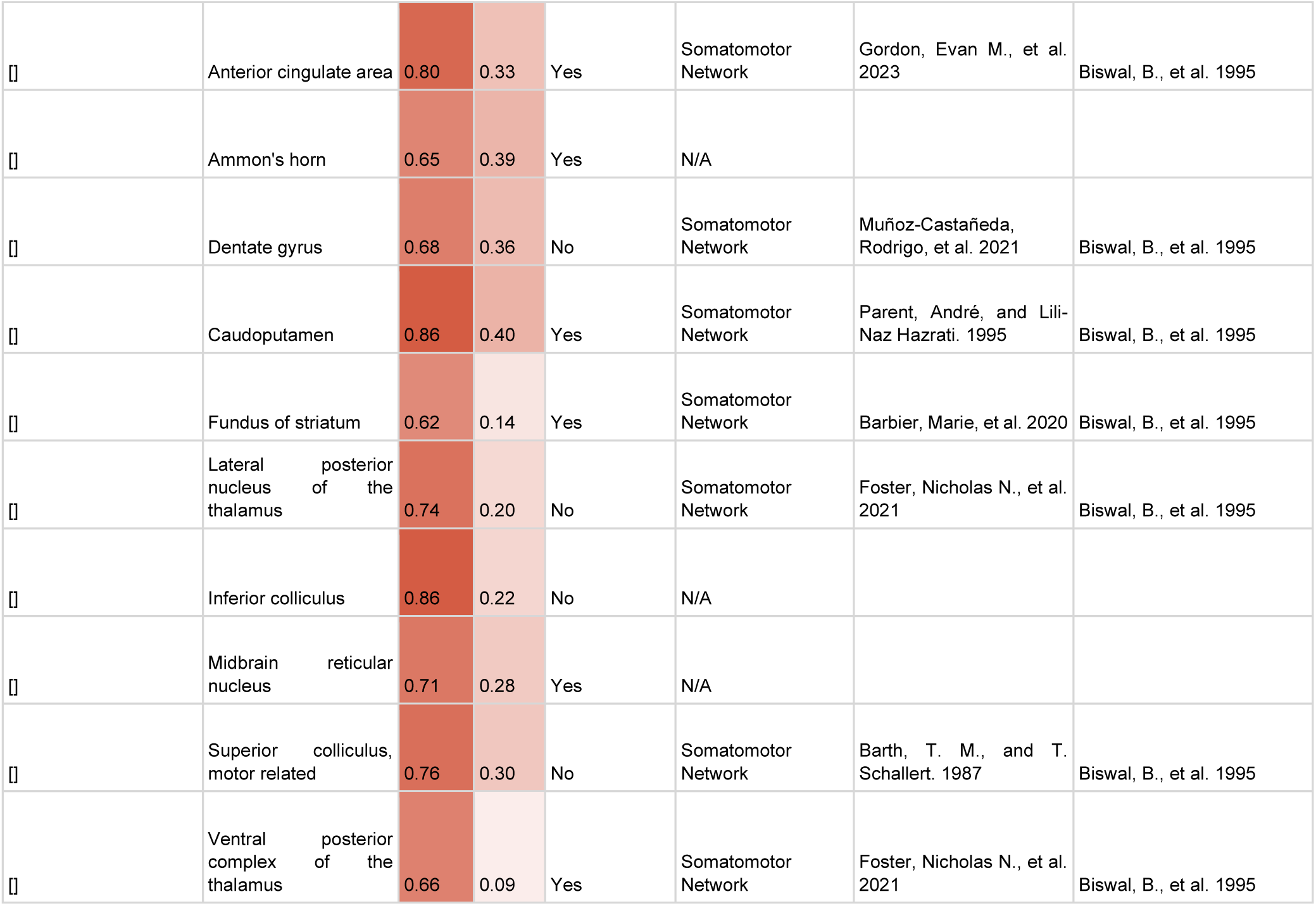
Literature Search Results. Column 1 = Seed ROI, Column 2 = Suspected Connected ROI, Column 3 = PAU Correlation Coefficient (Darker reds from Column 2-3 represent higher correlation), Column 4 = fMRI Correlation Coefficient, Column 5 = Existence of projection overlap from seed through/to suspected, Column 6 = Seed and Suspected Connection Shared Network, Column 7 = Reference for Suspected Connection Shared Network, Column 8 = Reference for Seed Shared Network. Blue highlighted rows represent connections not established by any of the three established routes.

## 4 Discussion

### 4.1 Mapping & Filtering

Pipelines exist to map and process fMRI mouse brain data, however a pipeline to map and process ultrasound and photoacoustic data for the purpose of whole-brain connectome generation has not previously been established. In this project we established that nearly the same pipeline used in fMRI mapping can be used to map and process PAU and ultrasound data; with the exception of temporal filtering and an additional step of converting exported.tif files to a single.nii file. This pipeline could be further streamlined for PAU and ultrasound data by extracting the SPM scripts used and implementing them into a single program. The segmentation process of the ultrasound data also has the potential to be automated if an appropriate artificial intelligence model was trained.

### 4.2 Functional Connectome

Our analysis of connectomes generated from fMRI compared to PAU revealed that the greatest similarities were present between the non-temporally filtered fMRI connectome and the PAU connectome. This suggests that, due to the inability to temporally filter the PAU data, the resulting connectome has similar false positives to the non-temporally filtered fMRI data that are present due to motion and low wavelength fluctuations. Additionally, the PAU connectome identified stronger and more functional connections. This difference may be due to the inherent nature of how oxygenation levels are detected in fMRI vs PAU. The fMRI only detects how the presence or absence of deoxyhemoglobin affects the MR signal, while PAU detects a signal change directly from deoxyhemoglobin and a separate signal change directly from oxyhemoglobin. The detection difference may give PAU a slight advantage in accuracy, which could account for the additional identified connections.

This fMRI mapping was limited in its physical range owing to the use of a RF surface coil which prevents analysis of more comprehensive brain activity. PAU also on the other hand does not require a surface coil for signal transmission and reception, and as such is not limited to acquisitions in those areas. The expanded view PAU offers allows for greater investigation into the brain networks inaccessible to many fMRI setups. The validation of these additional ROIs is crucial for confirming the accuracy and consistency of the PAU generated connectomes. This will be done in future research through additional literature review and alternate coil positions or improved coil ranges for fMRI acquisition.

The alternate parameter experiments revealed that, with the present technology, a step size of 0.5mm with low persists and both 750 nm and 850 nm emissions is able to provide optimal scans for connectome generation. The parameters that stand to see the greatest improvement when comparing PAU to fMRI is the sampling rate and temporal sample size. The difference in sampling rate between 1/42 Hz to 1 Hz prevented temporal filtering on the PAU data and introduced a large margin of error. The relevant brain activity frequencies for functional connectivity range from 0-0.1Hz (Cordes, Dietmar, et al. 2001). Thus, this ultrasound approach is only gathering the lower relevant frequencies and not the full range. If the sampling rate could be increased to a minimum of 1/5 Hz, all of the relevant frequencies could be adequately captured and temporal filtering could be performed. With an increase such as that, we would expect the elimination of many potential false positives resulting from motion or high frequency noise that could previously not be filtered.

### 4.3 Scientific Literature Review

Our literature review confirmed the existence of functional connectivity between many of the strong connections identified with PAU. Similar to the findings of Zhang et al. and Nasiriavanaki et al., our research strongly suggests that PAU is a viable approach for the generation of brain connectomes. The connections that could not be confirmed through the corresponding fMRI data, physical connections, or functional connectivity in literature suggest the possibility of previously unidentified connections or false positives. These connections could be further researched through experiments using fMRI with greater spatial resolution and increased sensitivity, additional PAU experiments with larger sample sizes, and PAU experiments with improved acquisition parameters such as faster sampling rate.

## 5 Conclusion

In conclusion, we have successfully established a pipeline for mapping and processing PAU data for the intended purpose of connectome generation similar to existing pipelines used for fMRI data. Our analysis demonstrated that the PAU generated connectomes, which modeled the entire brain, were comparable to fMRI connectomes. They most closely resembled connectomes derived from non-temporally filtered fMRI. That resemblance is indicative of similar false positives present in both data forms due to motion and low frequency noise that was not filtered out. PAU identified many connections that fMRI did not, and these connections were verified through literary research of several previously studied and recorded functional connections. This ability of PAU to detect connections established in literature that our fMRI data could not is likely due to an increased or alternate form of sensitivity via PAU’s dual detection of deoxyhemoglobin and oxyhemoglobin signals. Additionally, PAU’s lack of reliance on a surface coil enabled a broader physical range that can be used for more comprehensive brain connectome analysis. Future research of this modality as a viable alternative to fMRI should focus on the validation of the connections identified in PAU, the optimization and improvement of sampling rates, literature review of PAU connections beyond the physical range of our fMRI scan, and alternate experimental setups further comparing PAU and fMRI. Our findings strongly suggest that PAU is a viable alternative approach for generation of brain connectomes, with the potential to uncover previously unidentified functional connections and allow for more comprehensive brain analysis than traditional fMRI methodologies.

## Funding acknowledgements

State of Minnesota, Office of Higher Education, Spinal Cord and Traumatic Brain Injury Research Program, Department of Defense/Henry Jackson Foundation, the Suzanne M. Schwarz Fund and NIH funding R01 NS118330.

## Animal approval

All procedures on live animals were approved by the University of Minnesota’s Institutional Animal Care and Use Committee under protocol 2301–40677A.

## Software Links

SPM12: https://www.fil.ion.ucl.ac.uk/spm/software/spm12/ 3D Slicer: https://download.slicer.org/

Allen Brain Atlas nifti files: https://scalablebrainatlas.incf.org/mouse/ABA_v3

In-house Matlab scripts: https://github.com/icthreed/PAU-Connectome-Generation

## Supplemental data

Table 2: All acronyms and names of ROIs analyzed in adjacency matrices

